# Acetylcholine Signaling Genes are Required for Cocaine-Stimulated Egg Laying in *Caenorhabditis elegans*

**DOI:** 10.1101/2020.11.24.396333

**Authors:** Soren Emerson, Megan Hay, Mark Smith, Ricky Granger, David Blauch, Nicole Snyder, Rachid El Bejjani

## Abstract

The toxicity and addictive liability associated with cocaine abuse are well known. However, its mode of action is not completely understood, and effective pharmacotherapeutic interventions remain elusive. The cholinergic effects of cocaine on acetylcholine receptors, synthetic enzymes, and degradative enzymes have been the focus of relatively little empirical investigation. Due to its genetic tractability and anatomical simplicity, the egg laying circuit of the hermaphroditic nematode, *Caenorhabditis elegans*, is a powerful model system to precisely examine the genetic and molecular targets of cocaine *in vivo*. Here, we report a novel cocaine-induced behavioral phenotype in *Caenorhabditis elegans*, cocaine-stimulated egg laying. In addition, we present the results of an *in vivo* candidate suppression screen of synthetic enzymes, receptors, degradative enzymes, and downstream components of the intracellular signaling cascades of the main neurotransmitter systems that control *Caenorhabditis elegans* egg laying. Our results show that cocaine-stimulated egg laying is dependent on acetylcholine synthesis and synaptic release, functional nicotinic acetylcholine receptors, and the *Caenorhabditis elegans* acetylcholinesterases.

## INTRODUCTION

Cocaine abuse is estimated to account for 900,000 cases of substance use disorder and over 14,000 overdose deaths in the United States each year (Lipari and Van Horn 2013; Hedegaard et al. 2020). Despite the impact of cocaine abuse on public health, there are no FDA-approved pharmacotherapeutic interventions for either the addictive liability or toxicity associated with cocaine abuse (Kampman 2019). A major impediment to the development of effective pharmacological interventions is the non-specificity of the drug, as pharmacodynamic interventions designed to block the canonical monoamine neurotransmitters implicated in the mode of action of cocaine (dopamine (DA), serotonin (5-HT), and norepinephrine) have demonstrated limited efficacy in clinical trials (Pitts and Marwah 1989; Czoty et al. 2016). Elucidation of the full range of neurotransmitter systems and molecular effectors involved in the mode of action of cocaine could inform the development of novel pharmacotherapies.

Cholinergic signaling influences multiple processes underlying reward and dependence in the mammalian brain, including learning, attention, and motivation (Hasselmo 2006; Sofuoglu and Mooney 2009; Klinkenberg et al. 2011). In addition, behavioral and physiological research demonstrates an important role for acetylcholine (ACh) in the pathology of cocaine use disorder (Williams and Adinoff 2008). However, the cholinergic mechanisms of cocaine have not been fully investigated.

The *Caenorhabditis elegans* egg laying circuit has served as a tractable genetic and molecular model for the mechanistic underpinnings of various neurotransmitters, including ACh and the biogenic amines 5-HT, DA, octopamine (Oct), and tyramine (Tyr) (Schafer 2005; Chase 2007). The circuit consists of two hermaphrodite-specific motor neurons (HSNs) and six ventral type C motor neurons (VCs) that innervate eight vulval muscles (Schafer 2006). ACh plays a crucial and complex role in regulating the egg laying behavior of *C. elegans* (Rand 2007). The VCs, which are the primary cholinergic neurons in the egg laying circuit, innervate both the vulval muscles and the HSNs (White et al. 1986). Although typically regarded as serotonergic motor neurons, the HSNs have also been reported to function as cholinergic motor neurons in the egg laying circuit (Duerr et al. 2001; Pereira et al. 2015). ACh elicits excitatory and inhibitory effects on egg laying. The excitatory effect of ACh on egg laying is mediated by binding to nicotinic ACh receptors (nAChRs) on the vulval muscles inducing contraction and egg laying (Kim et al. 2001). Cholinergic inhibition of egg laying may result from binding of ACh to muscarinic ACh receptors (mAChRs) expressed in the egg laying circuit (Bany et al. 2003; Fernandez et al. 2020).

Here, we characterize a novel cocaine-induced behavioral phenotype in *C. elegans*, cocaine-stimulated egg laying, and present pharmacological and genetic evidence for the dependence of this behavioral phenotype on ACh, nAChRs, and acetylcholinesterases (AChEs). We show that the excitatory effect of cocaine on egg laying in *C. elegans* depends on genes involved in nicotinic cholinergic neurotransmission but not on the majority of genes encoding the molecular components of the other neurotransmitter systems expressed in the egg laying circuit. Together, our results provide a background for future investigation utilizing the *C. elegans* egg laying circuit to elucidate the cholinergic effects of cocaine.

## MATERIALS AND METHODS

### Nematode strains

The wildtype (WT) strain was Bristol N2. Animals were maintained on standard nematode growth media agar plates spread with *Escherichia coli* as a source of food (Stiernagle 2006). Temperature was controlled at 20 °C. Table S3 contains a complete list of all of the mutant *C. elegans* strains used in this study.

### Egg laying assays and statistical analysis

Egg laying assays in liquid media were performed according to a protocol by Moresco and Koelle (Hart 2006). For each experiment, an aqueous stock solution of 147.15 mM cocaine hydrochloride (50.000 mg/mL) was diluted to the desired treatment concentration in M9 buffer solution with an equal volume of 147.15 mM aqueous sucrose solution (50.369 mg/mL) diluted in M9 buffer solution in each of the control conditions. Late stage L4 hermaphrodites of the respective strains were picked ~24 hrs before assaying and continued to be cultured on agar plates with an OP50 lawn at 20°C. Ten day one adult hermaphrodites were isolated into single wells on a 96-well plate containing 35 μl experimental or equimolar control solutions for each replicate. After 1 hr, each well was examined, and the number of eggs laid by each animal was recorded.

For all liquid media egg laying assays, the investigators were blind to both the identity of the treatment and the genotype. Each experiment included 10 animals per experimental or control group, and each experiment was independently repeated five times. Seventeen total worms ruptured during experimental analysis. In these rare cases, the non-ruptured worms in the replicate were used to score the egg laying response.

For the analysis of the mean egg laying response, the mean number of eggs laid by each of the five replicate treatment groups was treated as a single data point. For the analysis of fraction responding, defined as the fraction of animals laying greater than or equal to one egg, the fraction responding of each of the five replicate treatment groups was treated as a single data point. Statistical analysis of the mean egg laying response and the fraction responding was performed using a Mann-Whitney U-test.

### Chemicals

Aqueous cocaine hydrochloride was supplied by the National Institute on Drug Abuse (Research Triangle Institute, Research Triangle Park, NC, USA).

### Data Availability Statement

All mutant strains examined in this study are available to order from the Caenorhabditis Genetics Center. The authors affirm that all data necessary for confirming the conclusions of this article are represented fully within the article and its tables and figures.

## RESULTS

### High concentrations of cocaine overcome the inhibition of egg laying in hypertonic liquid media

In an effort to add to the current cocaine abuse models available in *C. elegans*, we quantified any differences in egg laying in *C. elegans* treated with cocaine by assaying the mean egg laying response over a time period of 1 hr in hypertonic liquid media, a well-established method previously used to investigate the stimulation of egg laying by pharmacological agents (Horvitz et al. 1982; Trent et al. 1983; Desai and Horvitz 1989; Schafer and Kenyon 1995; Weinshenker et al. 1995; Jayanthi et al. 1998; Dempsey et al. 2005; Ward et al. 2009; Engleman et al. 2016; Engleman et al. 2018; Hart 2006). We tested the mean egg laying response of *C. elegans* to a series of increasing concentrations of cocaine (0 mM, 15.625 mM, 31.25 mM, and 62.5 mM) in liquid media for 1 hr (**Fig. 1A**). We show that treatment of WT *C. elegans* with 62.5 mM cocaine significantly stimulates egg laying compared to equimolar sucrose control [WT, 62.5 mM cocaine mean = 6.1±2.2; WT, 62.5 mM sucrose control mean = 0.34±0.56; *p* < 0.0001, N = 83 replicates (10 animals/replicate)] (**Fig. 1B**).

**Fig 1:**
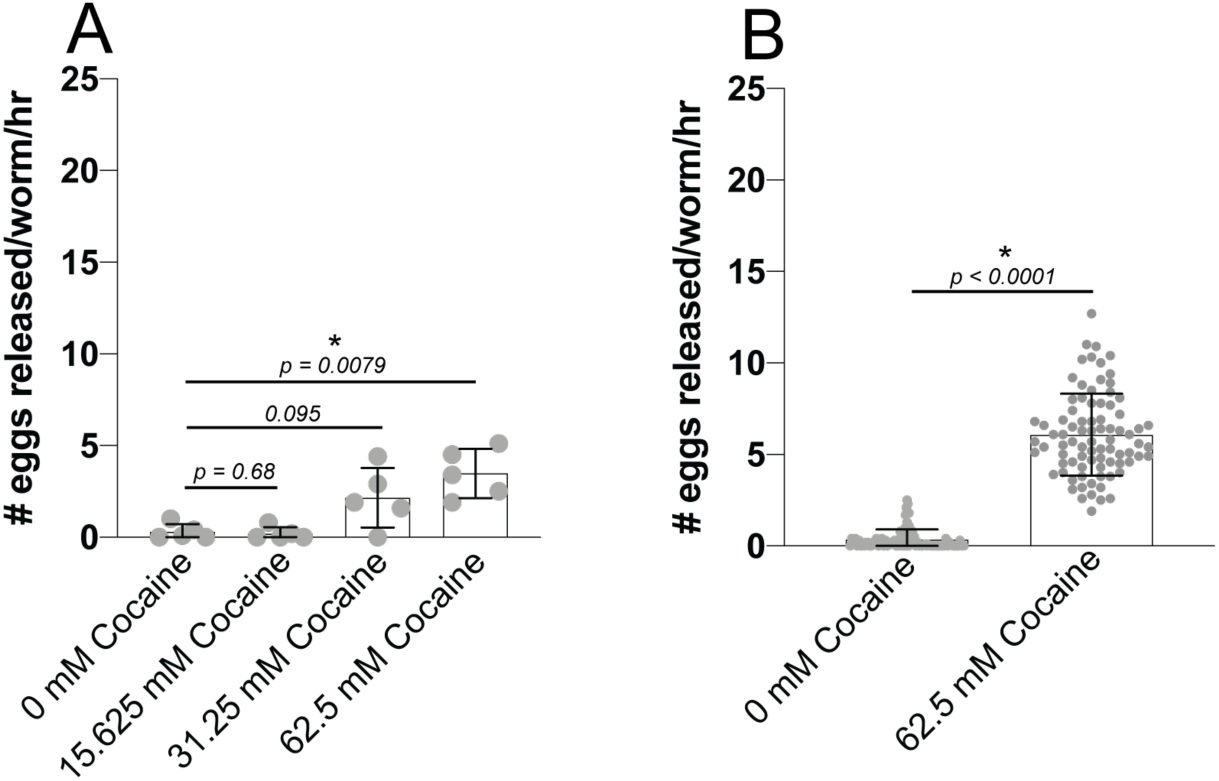
Cocaine stimulates *C. elegans* egg laying behavior. **A**. Egg laying response of *C. elegans* to a series of increasing concentrations of cocaine in liquid media. **B**. Egg laying response of *C. elegans* to 62.5 mM cocaine in liquid media compared to 62.5 mM sucrose control. For the data shown in A we performed five independent experiments with 10 animals in each group. The data shown in panel B show a compiled experiment for wild type worms from our experiments (83 independent experiments with 10 animals in each group). Error bars represent SD. Mann-Whitney U-test. Significance indicated by asterisks.

We used a 62.5 mM cocaine treatment and 62.5 mM sucrose control in all mutant analyses described in this paper. Interestingly, the concertation of cocaine necessary to robustly stimulate *C. elegans* egg laying behavior is greater than the concentrations of cocaine previously shown to modulate *C. elegans* locomotion by Ward *et al* (2009).

### ACh is the only neurotransmitter required for the stimulation of egg laying by cocaine in hypertonic liquid media

To begin investigating the neurobiological mechanism of cocaine-stimulated egg laying, we screened the egg laying response of individual mutants with defects in the synthetic enzymes for each of the neurotransmitters previously implicated in egg laying (**Fig. 2A**) (Schafer 2005). Interestingly, a reduction of function allele of *cha-1(p1152),* which encodes choline acetyltransferase (ChAT) necessary for ACh synthesis, suppresses egg laying in response to cocaine compared to WT [*cha-1(p1152)*, cocaine mean = 0.5±0.3; WT, cocaine mean = 4.8±1.8; *p* = 0.0079, N = 5 replicates (10 animals/replicate)] (**Fig. 2B**) (Rand and Russell 1984; Rand and Russell 1985).

**Fig 2:**
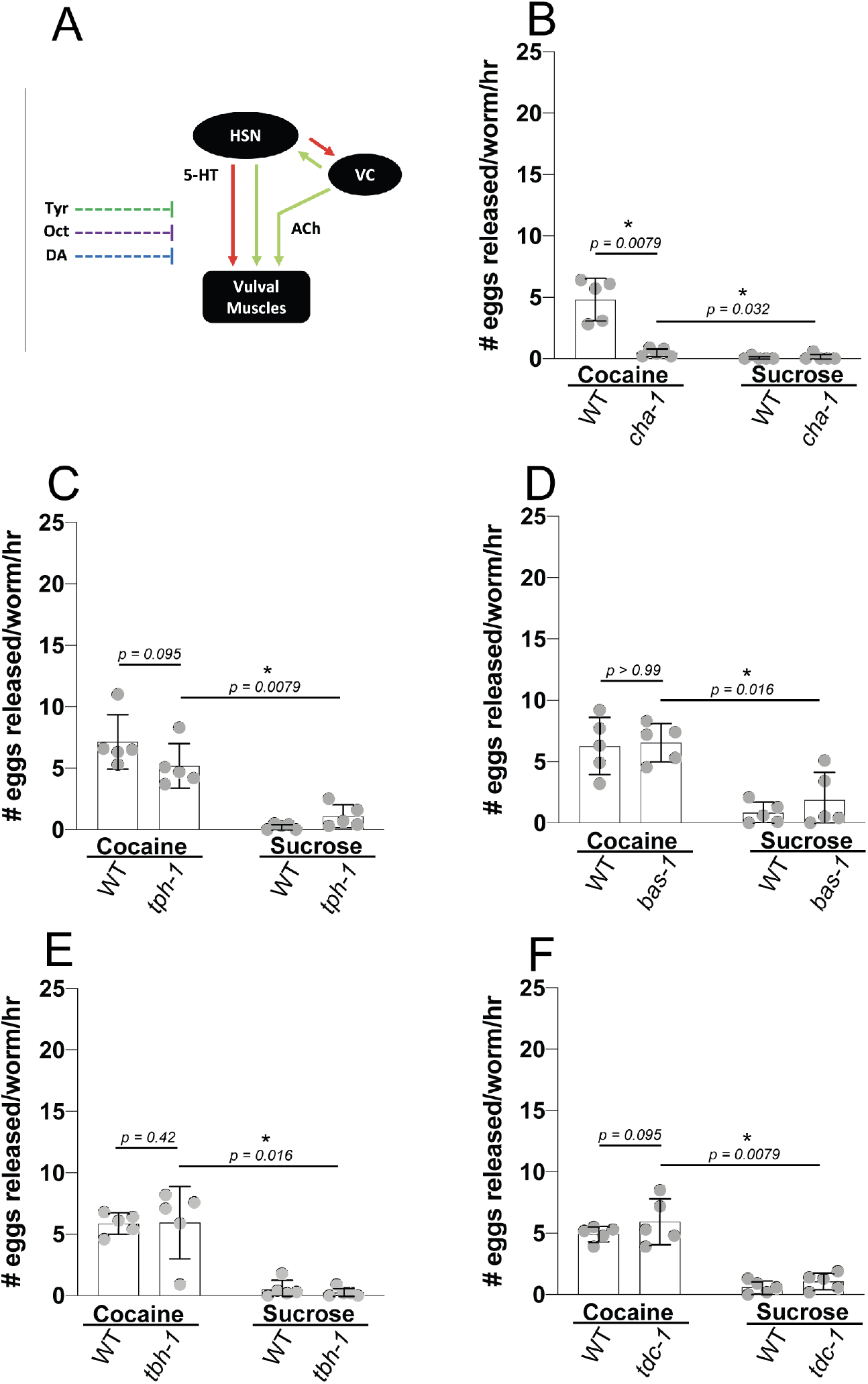
ChAT is required for the egg laying response to cocaine. **A**. Schematic of the *C. elegans* egg laying circuit showing HSNs, VCs, the vulval muscles, and the neurotransmitter systems expressed in the egg laying circuit, which include Tyr (dark green), Oct (purple), DA (blue), 5-HT (red), and ACh (light green). Extrasynaptic neurotransmitter systems marked with dashed lines. Synaptic neurotransmitter systems marked with solid lines. Egg laying behavior of mutants for **B**. ChAT/(*cha-1(p1152)*, **C**. TPH/*tph-1(mg280)*, **D**. AADC/*bas-1(ad446)*, **E**. TDC*/tdc-1(n3419)*, and **F**. TBH*/tbh-1(n3247)* in 62.5 mM cocaine or a 62.5 mM sucrose control compared to WT. We performed five independent experiments with 10 animals in each group. Error bars represent SD. Mann-Whitney U-test. Significance indicated by asterisks.

5-HT plays a well described excitatory role in the egg laying circuit, and 5-HT signaling is one of the canonical targets of cocaine in mammals (Higgins and Fletcher 2003; Schafer 2006; Müller et al. 2007; Schafer 2005; Chase 2007). Surprisingly, a putative null allele of the *C. elegans* tryptophan hydroxylase (TPH), the 5-HT synthetic enzyme, did not suppress cocaine-stimulated egg laying compared to WT [*tph-1(mg280)*, cocaine mean = 5.2±1.8; WT, cocaine mean = 7.1±2.2; *p* = 0.095, N = 5 replicates (10 animals/replicate)] (**Fig. 2C**) (Sze et al. 2000). These results suggest that 5-HT, one of the main modulators of egg laying behavior and a canonical cocaine effector in mammalian systems, is not a primary target of cocaine in this model system.

Additionally, putative null mutants for aromatic L-amino acid decarboxylase (AADC)/*bas-1(ad446)*, which encodes a 5-HT and DA synthetic enzyme, tyrosine beta-hydroxylase (TBH)*/tbh-1(n3247)*, which encodes an Oct synthetic enzyme, and tyrosine decarboxylase (TDC)/*tdc-1(n3419)*, which encodes a Tyr and Oct synthetic enzyme, do not exhibit a significant difference in the egg laying response to cocaine compared to WT, suggesting that neither DA, Tyr, nor Oct are necessary for cocaine-stimulated egg laying (**Fig. 2D-F**) (Hare and Loer 2004; Alkema et al. 2005).

### Cocaine-stimulated egg laying requires genes involved presynaptic ACh signaling

Suppression of the egg laying response to cocaine in reduction of function ChAT/*cha-1(p1152)* mutants suggests that presynaptic cholinergic neurotransmission is necessary for the egg laying response to the drug. To further examine this possibility, we assayed cocaine-stimulated egg laying in mutants with a reduction of function mutation in the vesicular ACh transporter (VAChT)/*unc-17(e245)* necessary for packaging of ACh within synaptic vesicles (Alfonso et al. 1993). We show that the egg laying response of VAChT/*unc-17(e245)* mutants is suppressed compared to WT [*unc-17(e245)*, cocaine mean = 0.5±0.1; WT, cocaine mean = 5.4±1.3; *p* = 0.0079, N = 5 replicates (10 animals/replicate)] and that cocaine does not stimulate a significant increase in egg laying in VAChT/*unc-17(e245)* mutants compared to equimolar sucrose control [*unc-17(e245)*, cocaine mean = 0.5±0.1; *unc-17(e245)*, sucrose control mean = 0.4±0.4; *p* = 0.64, N = 5 replicates (10 animals/replicate)] (**Fig. 3A**).

**Fig 3:**
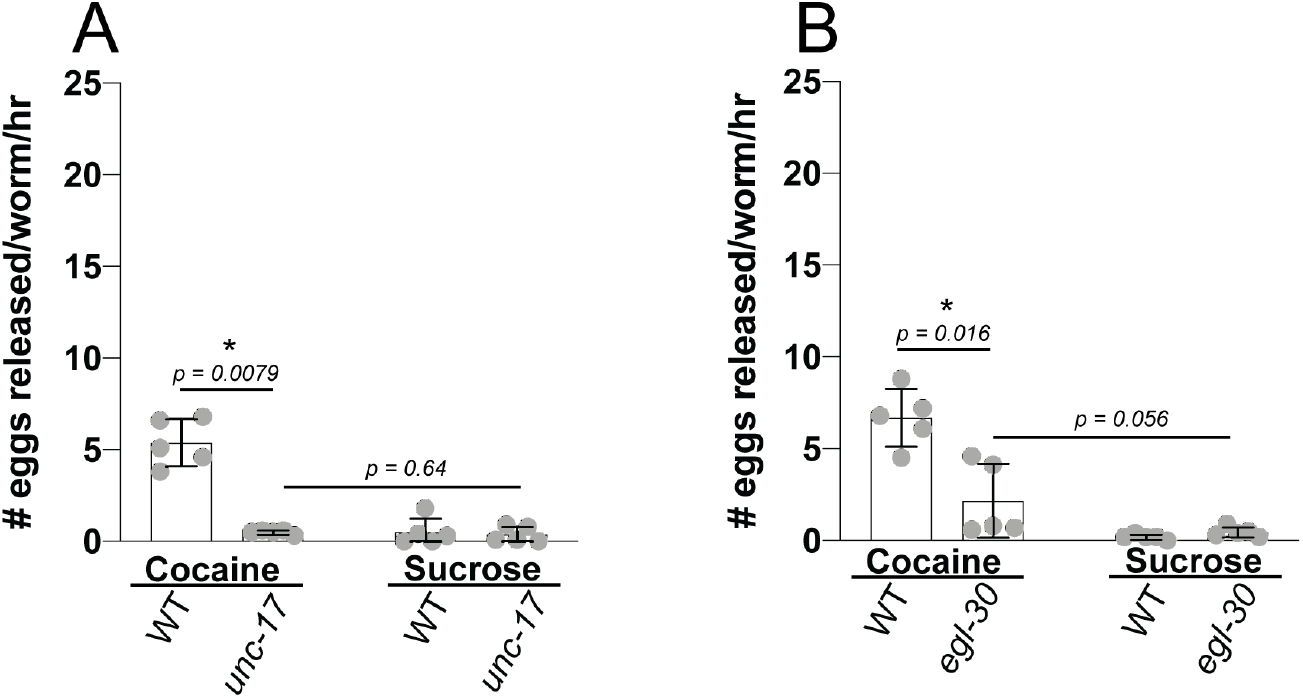
VAChT and Gα_q_ are required for the egg laying response to cocaine. Egg laying behavior of mutants for **A**. VAChT/*unc-17(e245)* and **B**. Gα_q_ *egl-30(ad806)*, which are required for ACh release from motor neurons, in 62.5 mM cocaine or a 62.5 mM sucrose control compared to WT. We performed five independent experiments with 10 animals in each group. Error bars represent SD. Mann-Whitney U-test. Significance indicated by asterisks.

The Gα_q_ ortholog encoded by *egl-30* in *C. elegans* is required for ACh release from motor neurons, among other neuronal functions (Lackner et al. 1999). We show that mutants with a reduction of function mutation in Gα_q_/*egl-30(ad806)* exhibit suppressed cocaine-stimulated egg laying compared to WT [*egl-30(ad806)*, cocaine mean = 2.2±2.0; WT, cocaine mean = 5.9±0.8; *p* = 0.016, N = 5 replicates (10 animals/replicate)] and that cocaine does not stimulate a significant increase in egg laying in Gα_q_/*egl-30(ad806)* mutants compared to equimolar sucrose control [*egl-30(ad806)*, cocaine mean = 2.2±2.0; *egl-30(ad806)*, sucrose control mean = 0.5±0.1; *p* = 0.056, N = 5 replicates (10 animals/replicate)] (**Fig. 3B**). Taken together, these results suggest that genes required for presynaptic ACh synthesis, packaging, and release are required for the egg laying response stimulated by cocaine in *C. elegans*.

### Genes encoding nAChRs, but not mAChRs, are required for the egg laying response to cocaine

To further examine the effect of cocaine on cholinergic signaling in the *C. elegans* egg laying circuit, we tested the egg laying response to cocaine in nAChR and mAChR mutants (**Fig. 4 and 5**) (Kim et al. 2001; Schafer 2006; Fernandez et al. 2020). Consistent with our finding that cholinergic neurotransmission is required for cocaine-stimulated egg laying, we show that the egg laying response to cocaine is significantly reduced compared to WT in *unc-29(e1072)* putative null mutants with defects in the beta subunit of nAChRs (CHRNB) [*unc-29(e1072)*, cocaine mean = 1.5±0.97; WT, cocaine mean = 6.0±1.7; *p* = 0.0079, N = 5 replicates (10 animals/replicate)] (**Fig. 4A**) (Fleming et al. 1997). Similarly, we show that the egg laying response to cocaine of alpha subunit nAChR putative null mutants (CHRNA)/*unc-38(e264)* is suppressed compared to WT [*unc-38(e264)*, cocaine mean = 3.8±1.2; WT, cocaine mean = 6.5±1.8; *p* = 0.024, N = 5 replicates (10 animals/replicate)] (**Fig. 4C**) (Richmond and Jorgensen 1999). These results suggest that nAChRs expressed on muscles, including the vulval muscles, are required for the molecular action of cocaine on *C. elegans* egg laying.

**Fig 4:**
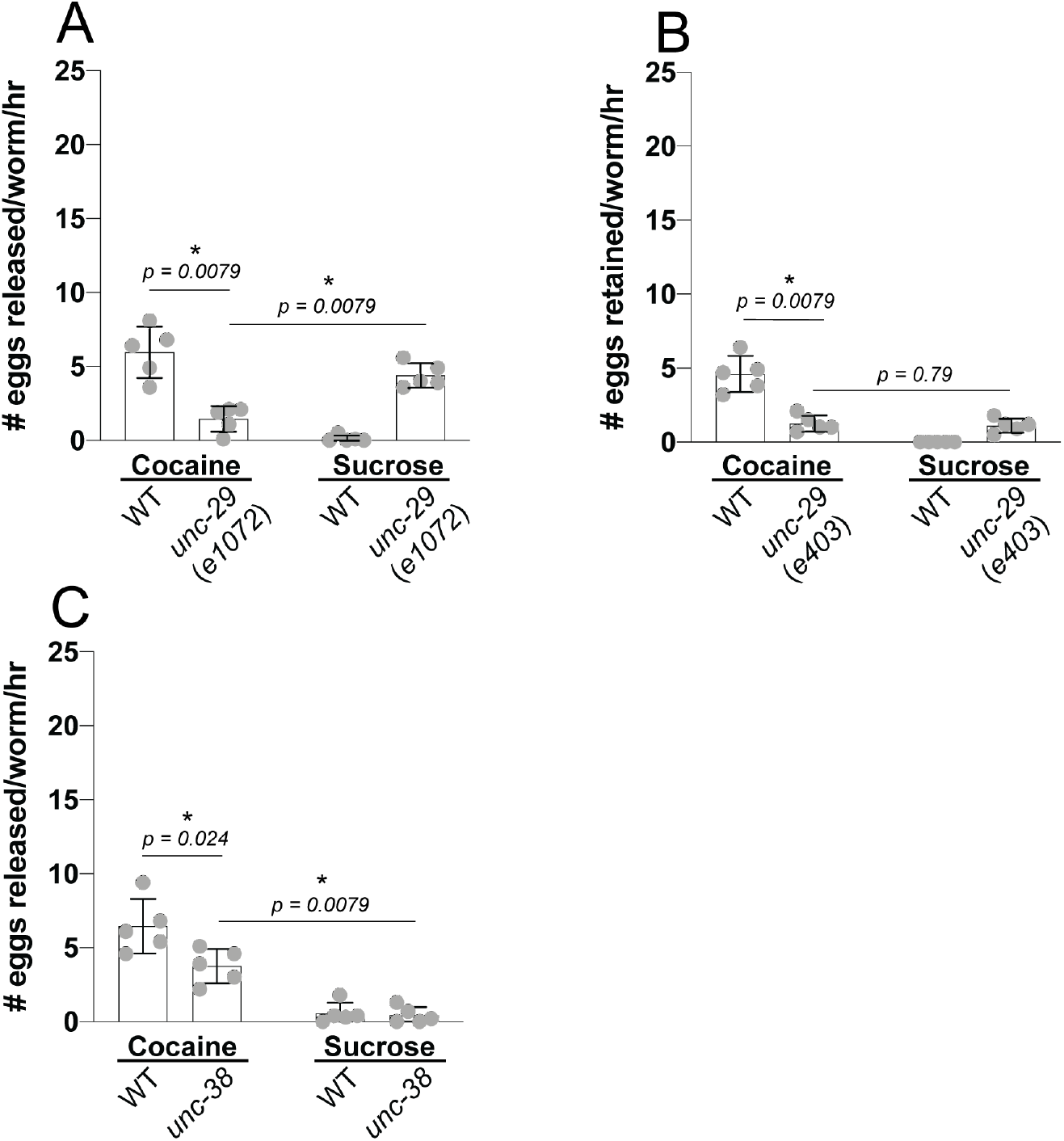
nAChRs are required for the egg laying response to cocaine. Egg laying behavior of **A**. CHRNB/*unc-29(e1072)*, **B**. CHRNB/*unc-29(e403)*, and **C**. CHRNA/*unc-38(e264)* in 62.5 mM cocaine or a 62.5 mM sucrose control compared to WT. We performed five independent experiments with 10 animals in each group. Error bars represent SD. Mann-Whitney U-test. Significance indicated by asterisks.

**Fig 5:**
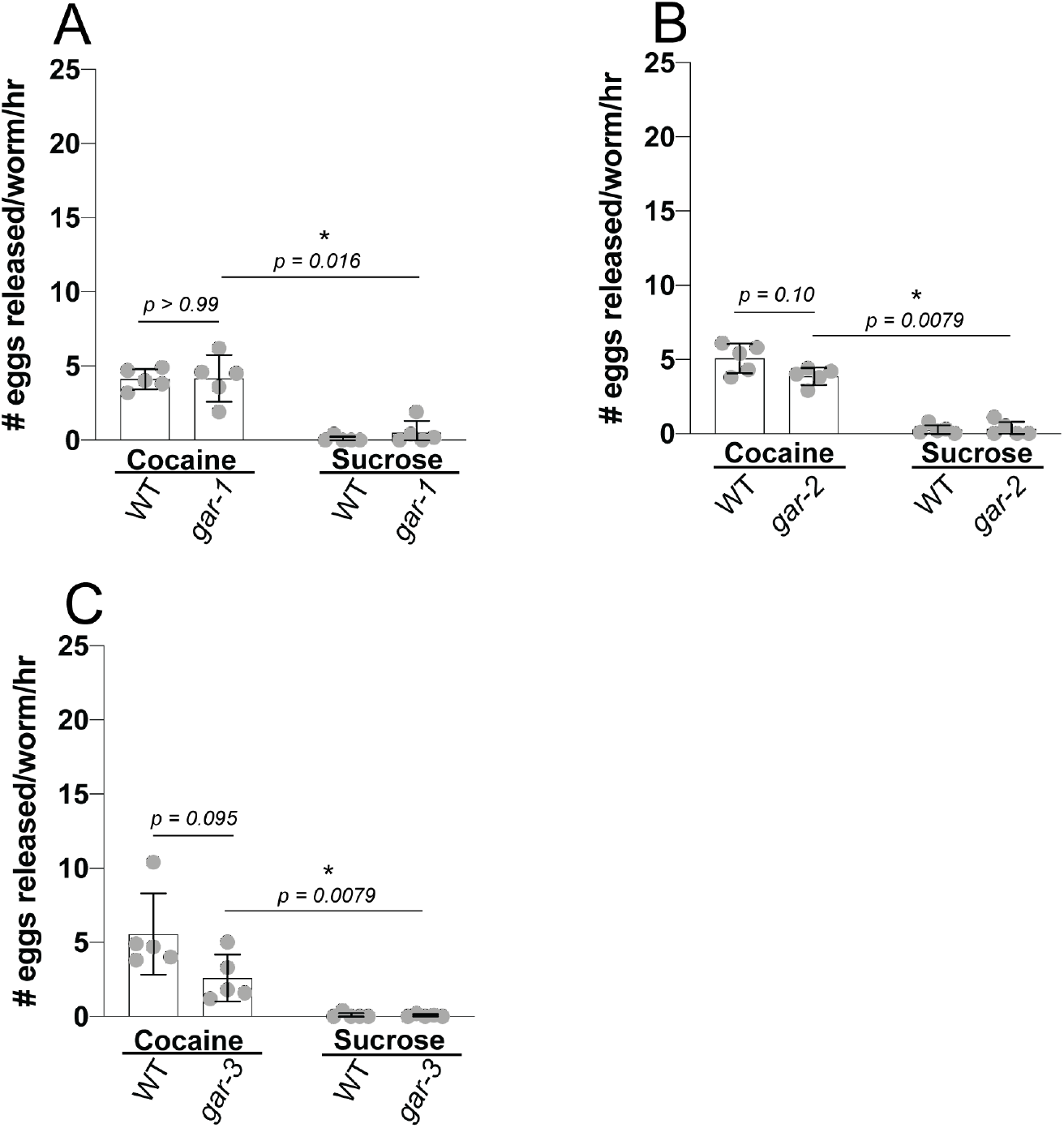
mAChRs are not required for the egg laying response to cocaine. Egg laying behavior of **A**. *gar-1(ok755)*, **B**. *gar-2(ok520)*, and **C**. *gar-3(gk305)* in 62.5 mM cocaine or a 62.5 mM sucrose control compared to WT. We performed five independent experiments with 10 animals in each group. Error bars represent SD. Mann-Whitney U-test. Significance indicated by asterisks.

The response to cocaine of CHRNB/*unc-29(e1072)* mutants revealed an unexpected result. Specifically, although cocaine-stimulated egg laying is suppressed in CHRNB/*unc-29(e1072)* mutants compared to WT animals treated with cocaine, CHRNB/*unc-29(e1072)* mutants treated with cocaine show a significantly reduced egg laying response compared to that of CHRNB/*unc-29(e1072)* mutants treated with equimolar sucrose control [*unc-29(e1072)*, cocaine mean = 1.5±0.97; *unc-29(e1072)* sucrose control mean = 4.4±0.8; *p* = 0.0079, N = 5 replicates (10 animals/replicate)] (**Fig. 4A**). To determine if this effect is generalizable for other alleles of CHRNB/*unc-29*, we repeated the experiment in a second CHRNB/*unc-29(e403)* allele. We show that, as observed for the CHRNB/*unc-29(e1072)* allele, CHRNB/*unc-29(e403)* mutants also exhibit a suppressed egg laying response to cocaine compared to WT [*unc-29(e403)*, cocaine mean = 1.3±0.6; WT, cocaine mean = 4.6±1.2; *p* = 0.0079, N = 5 replicates (10 animals/replicate)]. However, unlike CHRNB/*unc-29(e1072)*, the egg laying response to cocaine of CHRNB/*unc-29(e403)* mutants does not significantly differ from the egg laying response in CHRNB/*unc-29(e403)* treated with sucrose control [*unc-29(e403)*, cocaine mean = 1.3±0.6; *unc-29(e403)*, sucrose control mean = 1.1±0.5; *p* = 0.79, N = 5 replicates (10 animals/replicate)] (**Fig. 4B**). These results confirm the dependence of the egg laying response to cocaine on CHRNB/*unc-29* and suggest that the unexpected, reduced egg laying response to cocaine compared to sucrose control in CHRNB/*unc-29(e1072)* mutants may be allele-specific or due to an unidentified background mutation in this strain.

In addition to nAChRs, the *C. elegans* egg laying circuit expresses several mAChRs (Fernandez et al. 2020). To determine if mAChRs are required for the molecular mechanism of cocaine-stimulated egg laying, we assayed the egg laying response to cocaine of putative null mutants for the three mAChRs encoded by *gar-1(ok755)*, *gar-2(ok520)*, and *gar-3(gk305)* (Bany et al. 2003; Liu et al. 2007). We show all three mAChR mutants do not exhibit a significant difference in the egg laying response to cocaine compared to WT (**Fig. 5A-C**).

### AChE-encoding genes are required for the egg laying response to cocaine

AChEs catalyze the degradation of ACh at the synapse, decreasing cholinergic neurotransmission (Dvir et al. 2010). Biochemical analysis has revealed multiple genes encoding *C. elegans* AChEs, including *ace-1*, *ace-2*, *ace-3*, and *ace-4* (Combes et al. 2001). The AChEs encoded by *ace-1* and *ace-2* are the major degradative enzymes of ACh whereas the AChE encoded by *ace-3* accounts for a minor proportion of AChE activity, and the AChE encoded by *ace-4* is transcribed but does not result in a catalytically active protein (Johnson and Russell 1983; Kolson and Russell 1985; Arpagaus et al. 1998; Rand 2007). In order to explore all aspects of the relationship between cocaine and cholinergic signaling in the egg laying response, we tested the egg laying response to cocaine of *ace-2(g72)*; *ace-1(p1000)* and *ace-4 ace-3(dc2)* putative null double mutants (Culotti et al. 1981; D. Combes et al. 2000). *ace-2(g72)*; *ace-1(p1000)* double mutants exhibit a significant decrease in the egg laying response to cocaine compared to WT [*ace-2(g72)*; *ace-1(p1000)*, cocaine mean = 0.20±0.19; WT, cocaine mean = 5.2±2.6; *p* = 0.0079, N = 5 replicates (10 animals/replicate)] (**Fig. 6A**) whereas *ace-4 ace-3(dc2)* double mutants exhibit an egg laying response to cocaine that does not differ significantly compared to WT (**Fig. 6B**).

**Fig 6:**
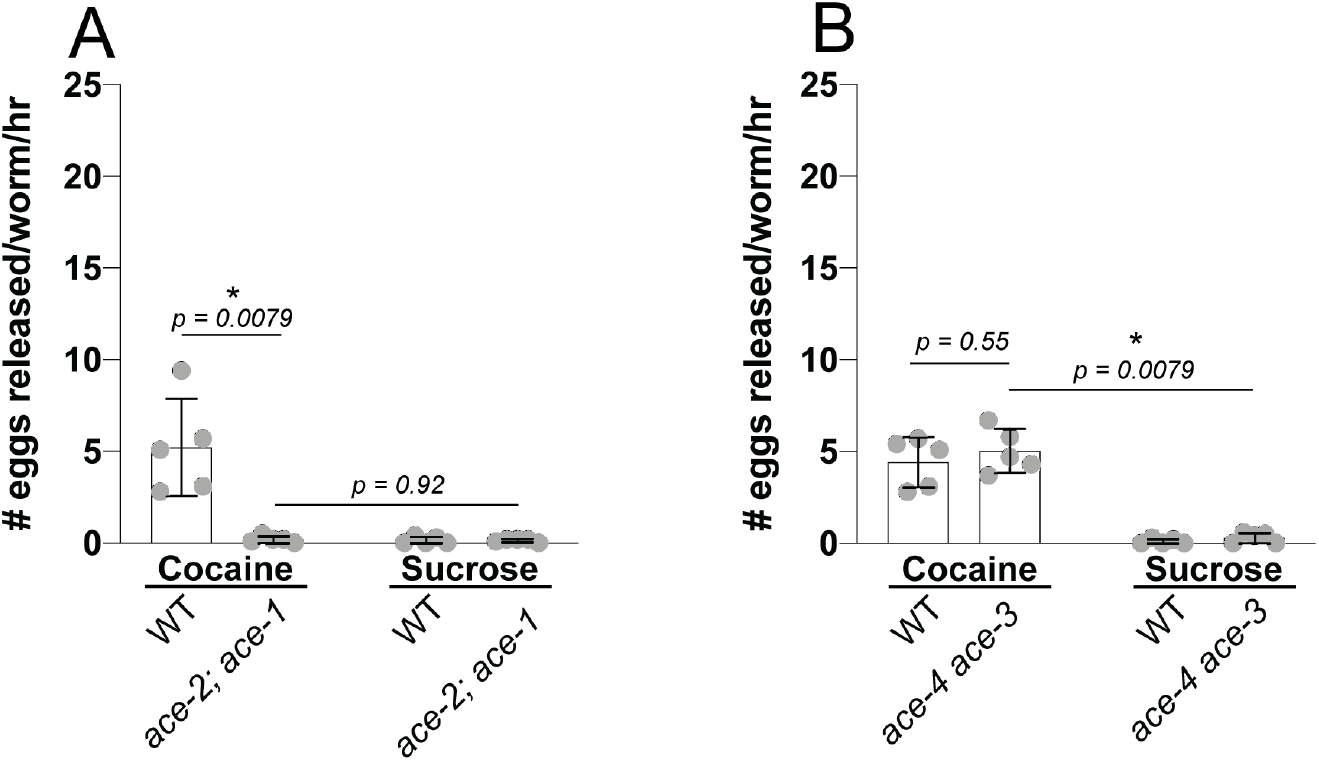
AChE-encoding genes are required for the egg laying response to cocaine. Egg laying behavior of **A**. *ace-2(g72)*; *ace-1(p1000)* and **B**. *ace-4 ace-3(dc2)* in 62.5 mM cocaine or a 62.5 mM sucrose control compared to WT. We performed five independent experiments with 10 animals in each group. Error bars represent SD. Mann-Whitney U-test. Significance indicated by asterisks.

### Cocaine-stimulated egg laying is dependent on the HSNs but not 5-HT receptor genes

The HSNs are the primary serotonergic motor neurons in the *C. elegans* egg laying circuit (Schafer 2006). In addition to their function as serotonergic motor neurons, the HSNs also function as cholinergic motor neurons (Duerr et al. 2001; Pereira et al. 2015). To further assess the role of the HSNs, 5-HT, and ACh in cocaine-stimulated egg laying, we quantified the egg laying response to cocaine in animals containing a semi-dominant allele of *egl-1*, shown to trigger HSN cell death (Conradt and Horvitz 1998). We show that *egl-1(n487)* mutants exhibit a slight but statistically significant difference in the egg laying response to cocaine compared to WT [*egl-1(n487)*, cocaine mean = 7.2±1.2; WT, cocaine mean = 9.3±0.9; *p* = 0.032, N = 5 replicates (10 animals/replicate)] (**Fig. 7A**). In addition, our data show that the HSNs contribute to cocaine-stimulated egg laying but that their presence is not necessary for the behavior. Specifically, cocaine stimulates a significant increase in egg laying compared to equimolar sucrose control in animals lacking the HSNs [*egl-1(n487)*, cocaine mean = 7.2±1.2; *egl-1*(*n487*), sucrose control mean = 1.2±0.9; *p* = 0.0079, N = 5 replicates (10 animals/replicate)].

**Fig 7:**
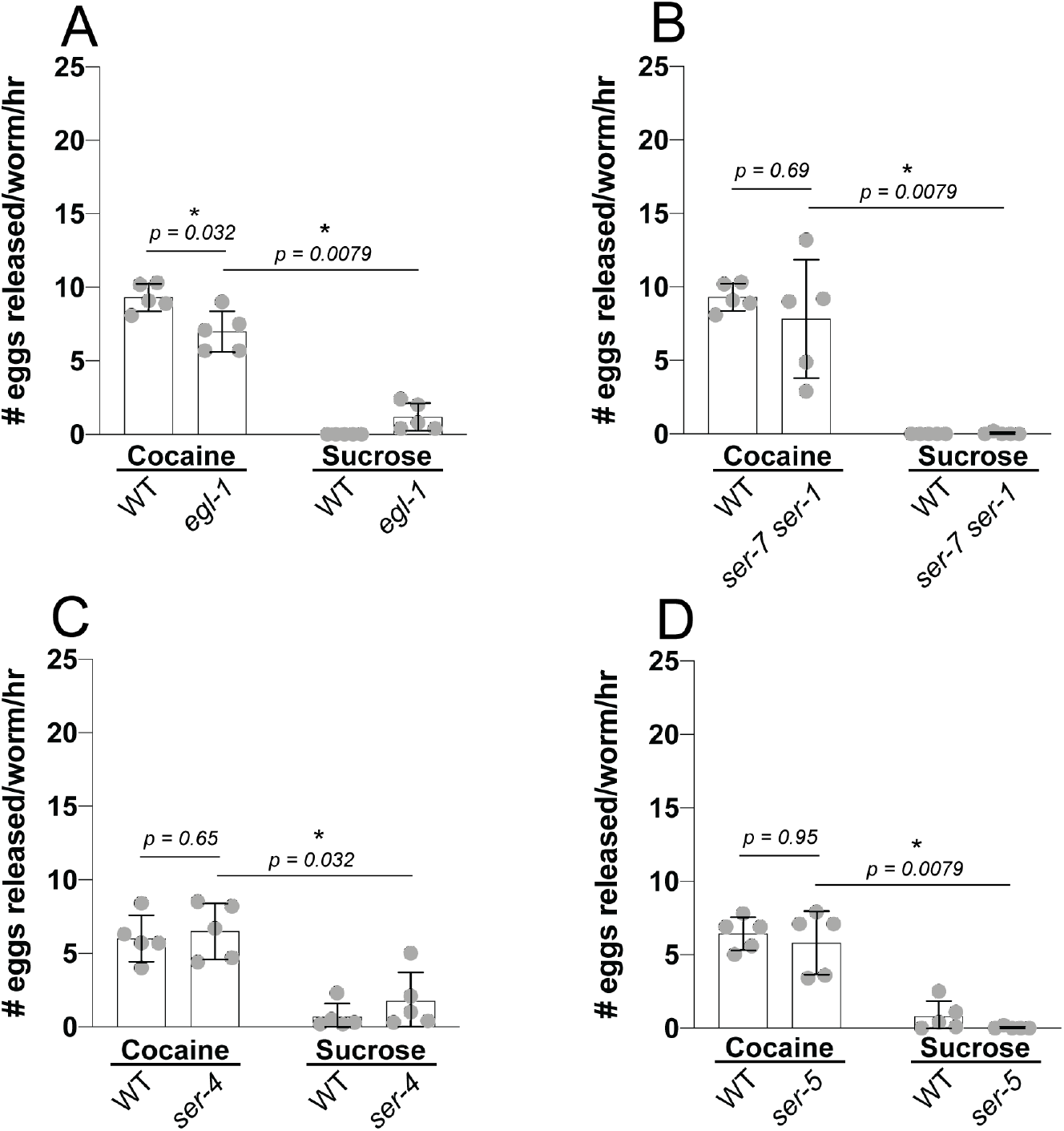
The HSNs, but not 5-HT receptors, are required for the egg laying response to cocaine. Egg laying behavior of **A.** animals lacking HSNs/*egl-1(n487)*, **B**. *ser-7(tm1325) ser-1(ok345)* double mutants, **C**. *ser-4(ok512)* mutants, and **D***. ser-5(ok3087)* mutants in 62.5 mM cocaine or a 62.5 mM sucrose control compared to WT. We performed five independent experiments with 10 animals in each group. Error bars represent SD. Mann-Whitney U-test. Significance indicated by asterisks.

Our previous analysis of mutants with defects in cholinergic and serotonergic neurotransmission indicates that ACh, but not 5-HT, is required for the stimulatory effects of cocaine on *C. elegans* egg laying behavior. To determine if cocaine acts on 5-HT receptors in the egg laying circuit, we tested the egg laying response to cocaine in putative null mutants with defects in four metabotropic 5-HT receptors expressed in the *C. elegans* egg laying circuit. The genotypes tested include *ser-5(ok3087)*, which encodes an excitatory Gα_s_-coupled 5-HT receptor, *ser-4(ok512),* which encodes an inhibitory Gα_o_-coupled 5-HT receptor, and a *ser-7(tm1325) ser-1(ok345)* double mutant, which encode an excitatory Gα_q_-coupled 5-HT receptor and an excitatory Gα_s_-coupled 5-HT receptor, respectively (Olde and McCombie 1997; Carnell et al. 2005; Hobson et al. 2006; Churgin et al. 2017). We show that *ser-7(tm1325) ser-1(ok345)* double mutants and *ser-5(ok3087)* mutants exhibit an egg laying response to cocaine that does not differ significantly compared to WT, suggesting that the excitatory post-synaptic effects of 5-HT are not required for cocaine-stimulated egg laying (**Fig. 7B and D**). Further, our data suggest that cocaine does not stimulate egg laying by blocking the inhibitory effects of *ser-4(ok512)* (**Fig. 7C**). Together, our data show that the egg laying response to cocaine depends on ACh receptors, but not 5-HT receptors, and suggest that suppression of the egg laying response to cocaine in *egl-1(n487)* mutants may result from a lack of presynaptic cholinergic neurotransmission by the HSN motor neurons.

### Cocaine-stimulated egg laying is increased in 5-HT-gated chloride channel/*mod-1* mutants and 5-HT reuptake transporter/*mod-5* mutants

Cocaine induces a decrease in locomotion speed in *C. elegans* that is dependent on the gene *mod-5*, which encodes the *C. elegans* 5-HT reuptake transporter (SERT), and the gene *mod-1*, which encodes an inhibitory 5-HT-gated chloride channel (Ward et al. 2009). We tested the egg laying response to cocaine in 5-HT-gated chloride channel/*mod-1(ok103)* putative null mutants and SERT/*mod-5(n3314)* putative null mutants to examine the role of these genes in cocaine-stimulated egg laying (Ranganathan et al. 2000; Ranganathan et al. 2001). 5-HT-gated chloride channel*/mod-1(ok103)* mutants exhibit a significant increase in the egg laying response to cocaine compared to WT [*mod-1(ok103),* cocaine mean = 18±3.2, WT, 62.5 mM cocaine mean = 9.8±2.0; *p* = 0.0079, N = 5 replicates (10 animals/replicate)] (**Fig. 8A**). Similarly, SERT/*mod-5(n3314)* mutants exhibit a significant increase in the egg laying response to cocaine compared to WT [*mod-5(n3314)*, cocaine mean = 14.8±1.3; WT, cocaine mean = 9.8±2.0; *p* = 0.0079, N = 5 replicates (10 animals/replicate)] (**Fig. 8B**). The increased egg laying responses to cocaine in 5-HT-gated chloride channel*/mod-1(ok103)* mutants and SERT/*mod-5(n3314)* mutants suggest that the removal of 5-HT-gated chloride channel inhibition of egg laying or a global increase in 5-HT, respectively, may potentiate the egg laying response to cocaine.

**Fig 8:**
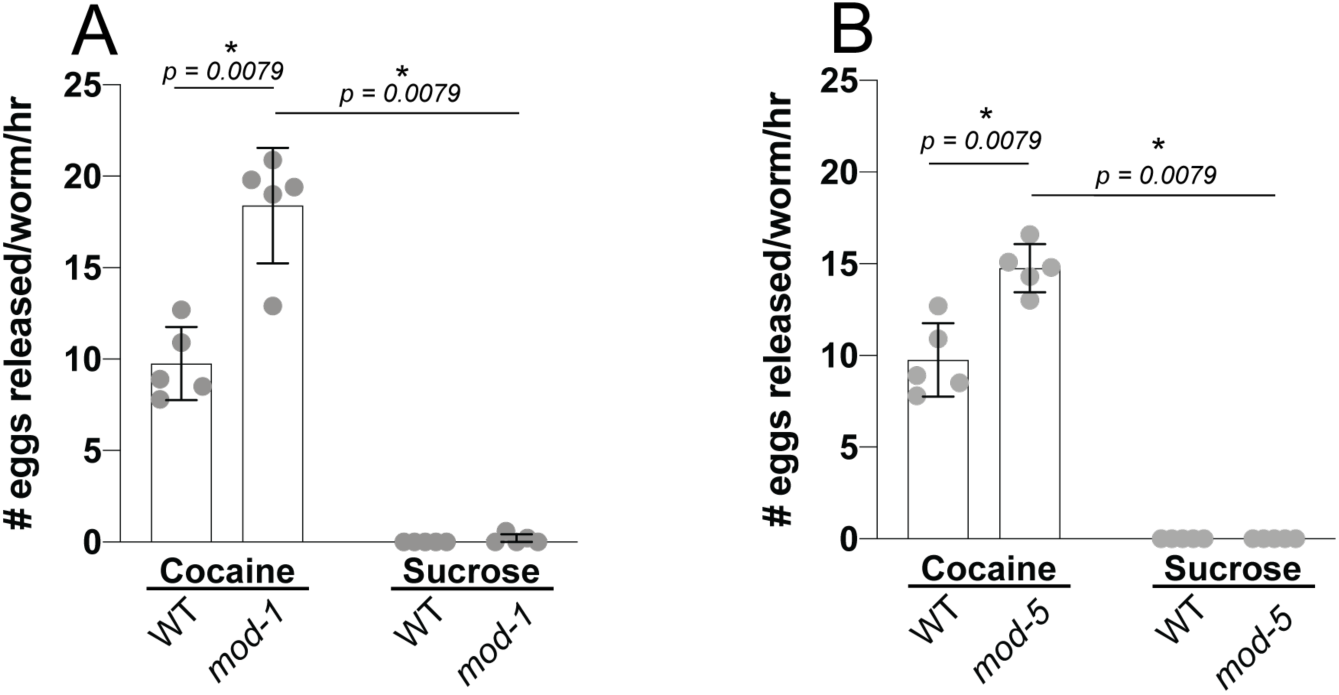
The mean egg laying response to cocaine is increased in 5-HT-gated chloride channel mutants and SERT mutants. **A.** 5-HT-gated chloride channel/*mod-1(ok103)* mutants, **B**. SERT/*mod-5(n3314)* mutants in 62.5 mM cocaine or a 62.5 mM sucrose control compared to WT. We performed five independent experiments with 10 animals in each group. Error bars represent SD. Mann-Whitney U-test. Significance indicated by asterisks.

### Cocaine-stimulated egg laying is weakly suppressed in DA reuptake transporter mutants but not mutants with defects in other DA signaling genes

The majority of mechanistic cocaine research has examined the drug’s activity as a DA reuptake inhibitor (Nestler 2005). According to this mechanism, cocaine binds to and blocks the DA reuptake transporter (DAT), preventing removal of DA from the synapse and thereby potentiating dopaminergic neurotransmission. *dat-1* encodes the *C. elegans* DAT, which has been shown to be sensitive to cocaine in heterologous systems (Jayanthi et al. 1998). To determine if DAT/*dat-1* is required for the egg laying response to cocaine in *C. elegans*, we tested the egg laying response to cocaine in DAT/*dat-1(ok157)* putative null mutants (McDonald et al. 2007). DAT/*dat-1(ok157)* putative null mutants exhibit a weak but significant decrease in the egg laying response to cocaine compared to WT [*dat-1(ok157)*, cocaine mean = 2.2±1.7, WT, cocaine mean = 5.7±1.2; *p* = 0.048, N = 5 replicates (10 animals/replicate)] (**Fig. 9A**). However, cocaine stimulates a significant increase in egg laying compared to equimolar sucrose control [*dat-1(ok157)*, cocaine mean = 2.2±1.7; *dat-1(ok157)*, sucrose control mean = 0.2±0.3; *p* = 0.0079, N = 5 replicates (10 animals/replicate)].

**Fig 9:**
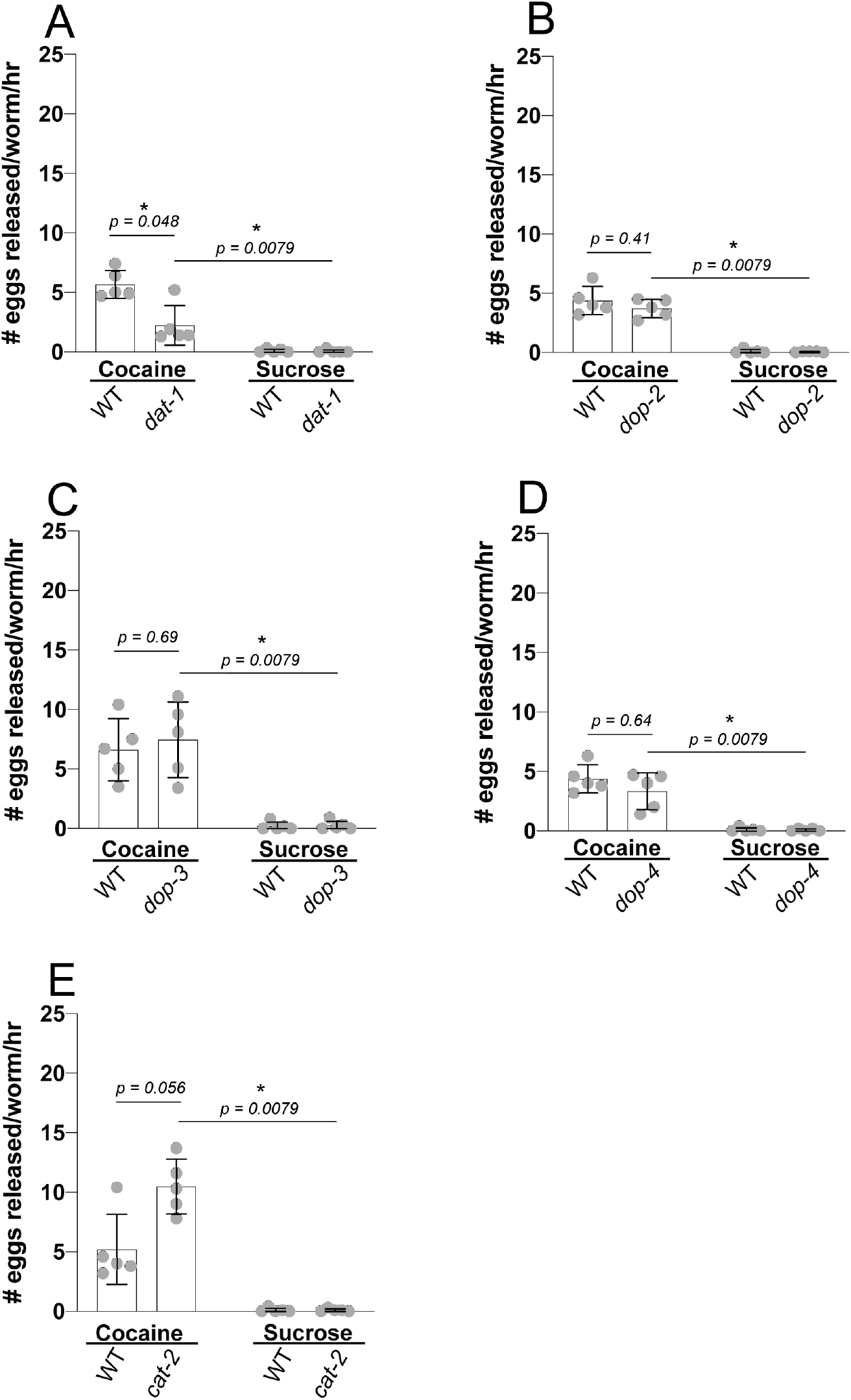
DAT is required for the egg laying response to cocaine, but not other DA signaling genes. Egg laying behavior of **A.** DAT mutants *dat-1(ok157)*, **B**. *dop-2(vs105)*, **C**. *dop-3(vs106)*, **D**. *dop-4(ok1321)*, and E. *cat-2(e1112)* in 62.5 mM cocaine or a 62.5 mM sucrose control compared to WT. We performed five independent experiments with 10 animals in each group. Error bars represent SD. Mann-Whitney U-test. Significance indicated by asterisks.

Our analysis of the egg laying response to cocaine in mutants for the DA and 5-HT synthetic enzyme, AADC/*bas-1(ad446)* (**Fig. 1D**), suggests that DA is not required for the egg laying response to cocaine. To further examine the role that DA plays in cocaine-stimulated egg laying, we tested the egg laying response to cocaine in a series of additional mutants with defects in DA signaling genes. The genotypes tested include putative null alleles of *dop-2(vs105)*, which encodes a Gα_o_-coupled DA receptor, *dop-3(vs106)*, which encodes a Gα_o_-coupled DA receptor, *dop-4(ok1321)*, which encodes a Gα_s_-coupled DA receptor, and *cat-2(e1112)*, which encodes the DA synthetic enzyme tyrosine hydroxylase (TH) (Lints and Emmons 1999; Chase et al. 2004; Cao and Aballay 2016; Fernandez et al. 2020). Consistent with our analysis of the egg laying response to cocaine in AADC/*bas-1(ad446)* mutants, neither TH/*cat-2(e1112)* mutants nor single mutants for the three genes encoding DA receptors expressed in the neurons and muscles that control *C. elegans* egg laying (*dop-2(vs105)*, *dop-3(vs106)*, and *dop-4(ok1321)*) exhibit a significant difference in the egg laying response to cocaine compared to WT (**Fig. 9B-E**). Together, these data suggest that DA is not a primary target of cocaine in this model system.

### Fraction of WT, cholinergic mutant, and monoamine mutant *C. elegans* that exhibit an egg laying response to cocaine

As is standard in the analysis of *C. elegans* egg laying behavior in response to pharmacological agents, we assayed the mean number of eggs laid in response to treatment by each of the genetic backgrounds tested compared to WT (Trent et al. 1983; Desai and Horvitz 1989; Schafer and Kenyon 1995; Weinshenker et al. 1995; Hart 2006). One limitation of comparing the mean eggs laid, however, is that it potentially obscures how the egg laying response to cocaine differs across the population of animals tested. To address this, we reanalyzed our data and calculated the fraction of wild type animals laying at least one egg in response to treatment with cocaine in hypertonic media for 1 hr (**Fig. 10A and B**). We show that cocaine stimulates an egg laying response in 85±15% of WT animals tested whereas only 14±16% of WT animals treated with equimolar sucrose control lay at least one egg [*p* < 0.0001, N = 83 replicates (10 animals/replicate)] (**Fig. 10B**).

**Fig 10:**
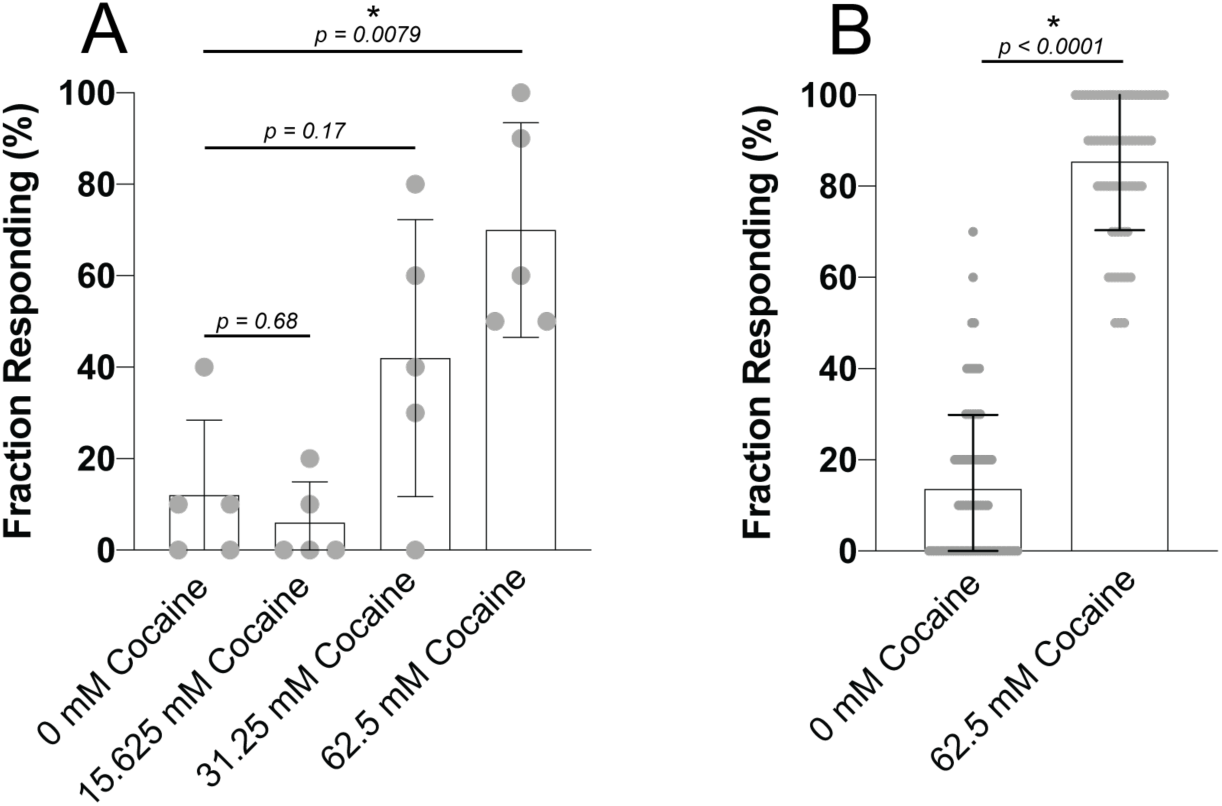
Fraction of WT *C. elegans* that emit an egg laying response to cocaine treatment. Fraction responding, defined as the fraction of animals releasing one or more eggs, of **A**. WT *C. elegans* to a series of increasing concentrations of cocaine in liquid media and **B**. WT *C. elegans* to 62.5 mM cocaine in liquid media compared to 62.5 mM sucrose control. For the data shown in A we performed five independent experiments with 10 animals in each group. For the data shown in panel B we re-analyzed 83 independent experiments with 10 animals in each group for which we had previously reported the mean number of eggs laid in 1 hr. Error bars represent SD. Mann-Whitney U-test. Significance indicated by asterisks.

We also reanalyzed the fraction responding to cocaine in all mutant strains tested compared to WT. The fraction of animals that lay at least one egg mostly followed the same pattern observed when analyzing the mean number of eggs laid in 1 hr (**Table. S1 and S2**).

## DISCUSSION

### Cocaine may modulate endogenous cholinergic neurotransmission to stimulate *C. elegans* egg laying

In the present study, we show that the mean egg laying response to cocaine in *C. elegans* is suppressed in mutants with defects in presynaptic cholinergic neurotransmission, postsynaptic defects in nAChRs, and defects in the *C. elegans* AChEs. Together, these results suggest that cocaine may modulate endogenous cholinergic neurotransmission to stimulate *C. elegans* egg laying through signaling at nAChRs. At this stage, the precise neurogenetic mechanism for the effects of cocaine on cholinergic neurotransmission remains unclear. Nonetheless, our novel findings demonstrate the utility of the *C. elegans* egg laying circuit as a model system for examination of the cholinergic effects of cocaine and lay the foundation for further mechanistic investigation.

A conclusive mechanism for regulation of cholinergic neurotransmission by cocaine will require the collection of additional data. To this end, there are several mechanistic investigations that we suggest may shed light on the cholinergic effects of cocaine in future studies.

Cocaine may act on an uncharacterized receptor to regulate cholinergic neurotransmission. A recent biochemical report identified brain acid soluble protein 1 as a novel, pharmacologically relevant receptor for cocaine in mammals (Harraz et al. 2020). Brain acid soluble protein 1 is not conserved in *C. elegans*, but the identification of a novel cocaine receptor illustrates the possibility that cocaine may act on novel, non-canonical targets. Future studies could utilize the genetic tractability of *C. elegans* to identify additional non-canonical cocaine targets that regulate cholinergic neurotransmission via forward genetic screens.

Additionally, cocaine may act simultaneously on one or more amine neurotransmitter systems or receptors to regulate cholinergic neurotransmission. *In vivo* micro dialysis studies in mammals have shown that by potentiating dopaminergic neurotransmission, cocaine increases ACh release from the nucleus accumbens and the hippocampus (Imperato et al. 1993; Mark et al. 1999). This raises the possibility that cocaine might act analogously through one or more amine neurotransmitters to regulate ACh release, possibly from the VCs, which are the primary cholinergic motor neurons in the *C*. *elegans* egg laying circuit or from the HSNs, which have also been reported to function as cholinergic motor neurons (Duerr et al. 2001; Schafer 2006; Pereira et al. 2015). Notably, none of the biogenic amine neurotransmitter mutants tested in the present study besides DAT/*dat-1* mutants exhibit a suppressed mean egg laying response to cocaine compared to WT. However, because the egg laying circuit expresses multiple functionally redundant receptors for each of the biogenic amine neurotransmitters, it is possible that our genetic analysis of single and double mutants was insufficient to suppress possible amine-dependent egg laying responses to cocaine (Fernandez et al. 2020). Future studies could examine the egg laying behavior of triple and quadruple mutants for the receptors and synthetic enzymes of ACh and each of the amine neurotransmitter systems expressed in the egg laying circuit.

Notably, in a model where the egg laying response to cocaine is dependent on increased ACh signaling, the suppression of the egg laying response to cocaine in *ace-2*; *ace-1* double mutants is unexpected. Interestingly, mammals express two distinct types of cholinesterases, a “true” AChE with high substrate specificity for ACh and a less substrate-specific “pseudo” butrylcholinesterase that is the primary metabolic enzyme of cocaine in mammals (Gorelick 1997; Masson and Lockridge 2010). The substrate specificity of AChEs in *C. elegans* is intermediate to the distinct mammalian AChE and butrylcholinesterase (Johnson and Russell 1983; Kolson and Russell 1985). This raises the possibility that cocaine may be a substrate for *C. elegans* AChEs. Consequently, the suppression of the egg laying response to cocaine in these mutants may indicate that *C. elegans* AChEs metabolize cocaine to a compound which stimulates egg laying through a nicotinic cholinergic pathway. Alternatively, if *C. elegans* AChEs metabolize cocaine, this may inhibit the enzymes’ function as ACh-degrading enzymes, causing an increase in ACh levels at nAChRs. The role that the *C. elegans* AChEs play in the mode of action of cocaine is intriguing firstly, because it may be indicative of a possible novel mechanism of cocaine action. Secondly, interest in developing recombinant metabolic enzymes of cocaine and its associated metabolites as pharmacotherapeutic interventions for cocaine overdose and cocaine use disorder has accelerated over the last decade (Zheng and Zhan 2008; Zhang et al. 2017). Additional biochemical and genetic investigation of the effect of cocaine on *C. elegans* AChEs may therefore be a fruitful area of future investigation.

### The increase in cocaine-stimulated egg laying in SERT/*mod-5* and 5-HT-gated chloride channel/*mod-1* null animals may be 5-HT independent

Our data show that animals with null mutations in SERT/*mod-5* and the 5-HT-gated chloride channel/*mod-1* exhibit an enhanced cocaine-stimulated egg laying response compared to WT animals. These data could be interpreted as aligned with a model in which cocaine increases 5-HT levels, inducing egg laying in WT animals, an effect that is expected to be further potentiated by a lack of 5-HT reuptake in SERT/*mod-5* null animals. This serotonergic mechanism is supported by a study that investigated a cocaine-dependent decrease in locomotion speed in *C. elegans*, which is suppressed in 5-HT-gated chloride channel/*mod-1*, SERT/*mod-5*, and the 5-HT synthetic enzyme, TPH/*tph-1* null mutants (Ward et al. 2009).

However, we show that the null mutant in the 5-HT synthetic enzyme, TPH*/tph-1(mg280)* and the double mutant in the 5-HT receptors, *ser-7(tm1325) ser-1(ok345),* do not suppress cocaine-stimulated egg laying, pointing to a 5-HT-independent mechanism in cocaine-stimulated egg laying. Additional data in SERT/*mod-5* and 5-HT signaling double mutants will be necessary to determine if the enhanced cocaine-stimulated egg laying observed in SERT/*mod-5* and 5-HT-gated chloride channel*/mod-1* null animals depends on endogenous 5-HT signaling.

Interestingly, recent reports using calcium imaging of the egg laying circuit in freely behaving worms show evidence of coordination between egg laying behavior, body bend, and locomotion (Zhang et al. 2008; Collins and Koelle 2013; Collins et al. 2016; Kopchock et al. 2021). Such detailed analysis of the coordinated function of both circuits in cocaine-treated WT and mutant animals *in vivo* will be highly valuable to directly describe the neurobiological mechanism of cocaine action in *C. elegans*.

### Potential of the egg laying circuit as a model system for the cholinergic effects of cocaine

In the present study, we show that, except for a putative null allele of DAT/*dat-1*, mutants with defects in the amine neurotransmitters DA, Tyr, Oct, and 5-HT do not exhibit a suppressed mean egg laying response to cocaine compared to WT. This is a surprising finding because cocaine’s canonical mechanism of action is potentiation of monoaminergic neurotransmission through blockade of monoamine reuptake transporters (Richards and Laurin 2020). We do, however, show that cocaine-stimulated egg laying is suppressed in animals with defects in nicotinic cholinergic neurotransmission, which suggests that cocaine may modulate endogenous ACh signaling to induce egg laying through stimulation of nAChRs on the vulval muscles. This possibility aligns with previous reports that cocaine increases ACh release in multiple structures in the limbic system (Imperato et al. 1993; Mark et al. 1999). At present, the mechanism underlying potential modulation of cholinergic neurotransmission by cocaine is unclear. Nonetheless, the results of this study provide a background for future investigation utilizing the *C. elegans* egg laying circuit to dissect the detailed molecular mechanisms underlying the potential conserved cholinergic effects of cocaine.

## Acknowledgments

This work was supported by NIH Grants DA045364, DA031725, and DA045714. The NIH had no role in the writing of the manuscript or in the decision to submit the manuscript for publication. Some strains were provided by the *Caenorhabditis* Genetics Center, which is funded by National Institutes of Health Office of Research Infrastructure Programs (P40 OD010440). Some strains were generated by the *C. elegans* Deletion Mutant Consortium. S.E. was supported by the R. Craig and Sheila Yoder Applied Research Fellowship (https://www.davidson.edu/academics/research/undergraduate-research/r-craig-and-sheila-yoder-applied-research-fellowship), the Davidson Research Initiative (https://www.davidson.edu/academics/research/undergraduate-research/davidson-research-initiative) and the Faculty Study and Research Grant from Davidson College (https://www.davidson.edu/academics/research/faculty-research/faculty-funding-opportunities-research). The authors have declared that no competing interests exist.

