## supplemental information for "Acetylcholine Signaling Genes are Required for Cocaine-Stimulated Egg Laying in *Caenorhabditis elegans*"

| Genetic Background | Defect | Fraction Responding ± SD (%) | N replicates (10 animals/replicate) | *p*-value |
|---|---|---|---|---|
| *dat-1(ok157)* | *C. elegans* DAT | 62±22 | 5 | 0.056 |
| WT control | | 96±5 | 5 | |
| *dop-2(vs105)* | Metabotropic DA receptor | 78±15 | 5 | >0.99 |
| WT control | | 78±16 | 5 | |
| *dop-3(vs106)* | Metabotropic DA receptor | 88±13 | 5 | 0.60 |
| WT control | | 94±9 | 5 | |
| *dop-4(ok1321)* | Metabotropic DA receptor | 78±22 | 5 | >0.99 |
| WT control | | 78±16 | 5 | |
| *cat-2(e1112)* | TH | 98±4 | 5 | 0.40 |
| WT control | | 88±16 | 5 | |
| *mod-1(ok103)* | 5-HT-Gated Chloride Channel | 100±0 | 5 | 0.44 |
| WT control | | 94±9 | 5 | |
| *mod-5(n3314)* | *C. Elegans* SERT | 96±5 | 5 | >0.99 |
| WT control | | 94±9 | 5 | |
| *egl-1(n487)* | Triggers HSN cell death | 92±9 | 5 | 0.44 |
| WT control | | 96±5 | 5 | |
| *ser-7(tm1325) ser-1(ok345)* | Metabotropic 5-HT receptors | 92±13 | 5 | 0.68 |
| WT control | | 96±5 | 5 | |
| *ser-5(ok3087)* | Metabotropic 5-HT receptor | 82±22 | 5 | >0.99 |
| WT control | | 84±13 | 5 | |
| *tph-1(mg280)* | TPH | 76±13 | 5 | 0.079 |
| WT control | | 92±8 | 5 | |
| *bas-1(ad446)* | AADC | 80±10 | 5 | 0.77 |
| WT control | | 82±15 | 5 | |

| | | | | |
|---|---|---|---|---|
| *ser-4(ok512)* | Metabotropic 5-HT receptor | 88±8 | 5 | >0.99 |
| WT Control | | 86±17 | 5 | |
| *tdc-1(n3419)* | TDC | 96±9 | 5 | 0.52 |
| WT control | | 90±10 | 5 | |
| *tbh-1(n3247)* | TBH | 98±5 | 5 | 0.72 |
| WT control | | 88±22 | 5 | |

**Table S1: Fraction of monoamine mutants that emit an egg laying response to cocaine treatment compared to WT.** Fraction responding, defined as the fraction of animals releasing one or more eggs, of monoamine mutants treated in 62.5 mM cocaine compared to WT. Error represented by SD. Mann-Whitney U-test. Significance indicated by asterisks.

| Genetic Background | Defect | Fraction Responding ± SD (%) | N replicates (10 animals/replicate) | *p*-value |
|---|---|---|---|---|
| *unc-38(e264)* | CHRNA | 80±12 | 5 | 0.53 |
| WT control | | 82±19 | 5 | |
| *ace-2(g72)*; *ace-1(p1000)* | AChEs | 10±7 | 5 | 0.0079* |
| WT control | | 86±11 | 5 | |
| *ace-4 ace-3(dc2)* | AChEs | 88±8 | 5 | >0.99 |
| WT control | | 88±13 | 5 | |
| *unc-29(e1072)* | CHRNB | 36±17 | 5 | 0.0079* |
| WT control | | 86±11 | 5 | |
| *unc-29(e403)* | CHRNB | 58±13 | 5 | 0.024* |
| WT control | | 88±16 | 5 | |
| *egl-30(ad806)* | Gαq | 43±32 | 5 | 0.0079* |
| WT control | | 90±10 | 5 | |
| *unc-17(e245)* | VAChT | 36±16 | 5 | 0.016* |
| WT control | | 74±15 | 5 | |
| *cha-1(p1152)* | ChAT | 36±21 | 5 | 0.0079* |
| WT control | | 92±13 | 5 | |
| *gar-2(ok520)* | mAChR | 84±12 | 5 | 0.27 |
| WT control | | 92±11 | 5 | |
| *gar-1(ok755)* | mAChR | 78±16 | 5 | 0.29 |
| WT control | | 86±15 | 5 | |
| *gar-3(gk305)* | mAChR | 62±18 | 5 | 0.032* |
| WT control | | 94±5 | 5 | |

**Table S2: Fraction of cholinergic mutants that emit an egg laying response to cocaine treatment compared to WT.** Fraction responding, defined as the fraction of animals releasing one or more eggs, of cholinergic mutants treated in 62.5 mM cocaine compared to WT. Error represented by SD. Mann-Whitney U-test. Significance indicated by asterisks.

| Strain Name | Genotype and allele designation | Mutation | Reference |
|---|---|---|---|
| MT15434 | *tph-1(mg280)* | Large Deletion; Putative Null | (Sze et al. 2000) |
| MT13113 | *tdc-1(n3419)* | Large Deletion; Putative Null | (Alkema et al. 2005) |
| MT9455 | *tbh-1(n3247)* | Large Deletion; Putative Null | (Alkema et al. 2005) |
| MT7988 | *bas-1(ad446)* | Large Deletion; Putative Null | (Hare and Loer 2004) |
| PR1152 | *cha-1(p1152)* | Substitution; Reduction of Function | (Rand and Russell 1984) |
| CB933 | *unc-17(e245)* | Substitution; Reduction of Function | (Alfonso et al. 1993) |
| CB1072 | *unc-29(e1072)* | Uncurated; Putative Null | (Fleming et al. 1997) |
| CB403 | *unc-29(e403)* | Substitution; Putative Null | (Gottschling et al. 2017) |
| CB904 | *unc-38(e264)* | Substitution; Putative Null | (Richmond and Jorgensen 1999) |
| RB896 | *gar-1(ok755)* | Large Deletion; Putative Null | (The C. elegans Deletion Mutant Consortium 2012) |
| RB756 | *gar-2(ok520)* | Large Deletion; Putative Null | (Bany et al. 2003; The C. elegans Deletion Mutant Consortium 2012) |
| VC657 | *gar-3(gk305)* | Large Deletion; Putative Null | (Liu et al. 2007) |
| GG201 | *ace-2(g72)*; *ace-1(p1000)* | Substitutions; Putative Null | (Culotti et al. 1981) |
| PR1300 | *ace-4 ace-3(dc2)* | Large Deletion; Putative Null | (Didier Combes et al. 2000) |
| MT1082 | *egl-1(n487)* | Gain of Function | (Conradt and Horvitz 1998) |
| DA1084 | *egl-30(ad806)* | Substitution; Reduction of Function | (Lackner et al. 1999) |

| | | | |
|---|---|---|---|
| DA2109 | *ser-7(tm1325) ser-1(ok345)* | Large deletion; Putative Null | (Carnell et al. 2005; Hobson et al. 2006) |
| MT9668 | *mod-1(ok103)* | Large deletion; Putative Null | (Ranganathan et al. 2000) |
| MT9772 | *mod-5(n3314)* | Large deletion; Putative Null | (Ranganathan et al. 2001) |
| RB2277 | *ser-5(ok3087)* | Large deletion; Putative Null | (Churgin et al. 2017) |
| AQ866 | *ser-4(ok512)* | Large deletion; Putative Null | (Hapiak et al. 2009; The C. elegans Deletion Mutant Consortium 2012) |
| RM2702 | *dat-1(ok157)* | Large deletion; Putative Null | (McDonald et al. 2007; The C. elegans Deletion Mutant Consortium 2012) |
| LX702 | *dop-2(vs105)* | Large deletion; Putative Null | (Chase et al. 2004) |
| LX703 | *dop-3(vs106)* | Large deletion; Putative Null | (Chase et al. 2004) |
| RB1254 | *dop-4(ok1321)* | Large deletion; Putative Null | (The C. elegans Deletion Mutant Consortium 2012; Cao and Aballay 2016) |
| CB1112 | *cat-2(e1112)* | Substitution; Putative Null | (Lints and Emmons 1999) |

**Table S3: *C. elegans* mutants and strains used in this work.**
